# Genetic Influence on the Sleep EEG is Distributed Across Multiple Traits

**DOI:** 10.64898/2026.11.26.769206

**Authors:** Sophie R. Saremsky, Shuqiang Chen, Bethany Staley, Allan I. Pack, Shaun Purcell, Michael J. Prerau

## Abstract

Sleep spindles are among the most promising neurophysiological biomarkers in sleep research: they support memory consolidation, track healthy aging, are altered in psychiatric, neurodevelopmental, and neurodegenerative disorders, and are highly trait-like, varying between individuals while remaining stable within them across nights. Twin studies have shown that conventional spindle features such as density, amplitude, and duration are heritable, but important aspects of spindle activity remain unstudied, including spindle coupling to the slow-oscillation phase, which supports memory consolidation, and the individualized timing patterns that dominate spindle production. Spindles are also only a subset of a broader class of transient oscillations spanning 2–16 Hz that are more stable across nights than conventionally detected spindles and are altered in disorders such as schizophrenia, and whose genetic basis is unknown. Addressing these gaps is difficult because twin cohorts are small, which limits the detection of modest genetic effects one trait at a time. We analyzed sleep EEG (C3/C4) from 47 monozygotic (MZ) and 32 dizygotic (DZ) same-sex twin pairs (mean age 26.9±6.4 years). The DYNAM-O toolbox characterized transient oscillatory peaks in N2/N3 sleep, yielding 32 traits: event features in four frequency bands, fast-spindle timing patterns, and relative spectral power. MZ–DZ differences in intraclass correlation (ICC) were tested trait by trait with permutation tests (100,000 permutations, false discovery rate (FDR) correction) and jointly with group tests and multivariate co-twin distance. Fast-spindle (11–16 Hz) duration (ICC: MZ 0.82 vs DZ 0.31), prominence (0.81 vs 0.38), and slow-oscillation power (0.55 vs −0.10) showed greater MZ than DZ resemblance, as did the previously unstudied fast-spindle refractory period (0.68 vs 0.16) and the slow-oscillation power of 9–11 Hz and 5–9 Hz events (all FDR-adjusted p≤0.048). Preferred slow-oscillation phase showed no MZ–DZ difference in any band. Joint tests were more sensitive than any single trait: across all 32 traits, MZ co-twins were closer than DZ co-twins (0.86 vs 1.14, p=0.00002) and the mean-Z test was highly significant (p=0.00009), whereas the strongest single trait, corrected for the number of traits tested, was not (p=0.10). Genetic influence on sleep EEG therefore extends beyond fast spindles to lower-frequency transient oscillations and spindle timing, and is distributed across many traits in a pattern that single-trait analyses understate, supporting multivariate sleep biomarkers.

## INTRODUCTION

Sleep spindles are brief bursts of ∼12–16 Hz neural oscillatory activity during non-rapid eye movement (NREM) sleep that have been closely linked to memory consolidation and aging (1–4). Alterations in spindles, including their density, amplitude, and duration have been reported in many disorders including Alzheimer’s disease, schizophrenia and autism (2, 5–7). Notably, spindle features are highly trait-like, varying substantially between individuals and remaining stable within individuals across nights (8–10). The remarkable stability and individual variability of spindle characteristics raise a fundamental question regarding the extent to which these individual differences are genetically influenced. Therefore, understanding the genetic contributions to spindle features is crucial for their potential use as biomarkers of aging and disease.

Twin studies provide a powerful framework for investigating this question by comparing the resemblance of monozygotic (MZ) and dizygotic (DZ) twins, who share approximately 100% and 50% of their segregating genetic variants, respectively. Previous studies have demonstrated heritability of sigma-band power, which covers the frequency range of fast and slow spindles (11, 12). Analyses of detected spindle events have also suggested heritable variation in spindle characteristics, including amplitude, density, duration, and frequency and genetic contribution may be robust to detection algorithms (13, 14). Together, these studies provide evidence for potential genetic contributions to individual differences in spindle characteristics.

While sleep twin studies are an effective methodology for evaluating the genetic contribution to sleep traits, enrollment challenges and high operational costs make these difficult studies to undertake and thus have produced studies with small sample sizes. Moreover, because the unit of measurement is the subject pair, the effective sample size is reduced by half relative to the number of enrolled individuals. Consequently, these factors severely limit the ability of current twin studies to identify heritable traits with statistical confidence within standard heritability models, potentially precluding the detection of weakly heritable traits entirely.

Furthermore, given the advancement of the sleep field over time, there exist some important features of the sleep EEG relating to sleep spindles for which the genetic contribution has yet to be explored. In particular, there has been, to our knowledge, no study of the heritability of spindle coupling with the phase of slow oscillations (SOs), which plays an important role in memory consolidation and changes with aging (3, 15–17). Additionally, recent work (10) has shown that spindle occurrence is primarily driven by its preceding event history, such that the timing of one spindle greatly influences the timing of the next spindle. These influences create highly individualized spindle timing patterns that are stable across nights and change with age. Despite the established genetic influences on conventional spindle characteristics, whether these distinct dimensions of temporal organization are similarly heritable remains unstudied.

Moreover, studies suggested that conventionally defined spindles represent only a subset of a broader class of transient events with spindle-like morphology (18). Traditional spindle detection methods have been shown to ignore a substantial proportion of sigma-band oscillatory events, and these broader events exhibit greater night-to-night stability than conventionally detected spindles. Extending beyond sigma band, researchers further showed the existence of multiple classes of spindle-like events across a much wider frequency range and across aging (8, 17). Importantly, alterations in these other transient oscillations have also been observed across a variety of clinical conditions. For example, increased low-alpha (7–10 Hz) and decreased theta (4–6 Hz) transient oscillatory activity has been reported in schizophrenia compared to healthy controls (8), as well as increased theta and low-alpha activity in patients with fibromyalgia (19). Therefore, these findings highlight a major gap in our understanding of the genetic contributions to these important yet largely unexplored features of broad oscillatory activity that are closely associated with aging and disorders.

In this study, we aim to characterize the extent to which individual differences in transient sleep oscillations reflect genetic influence and whether this influence is distributed across a broader range of features beyond traditionally studied spindle characteristics. We analyze 47 MZ and 32 DZ same-sex twin pairs (of 53 MZ and 35 DZ pairs recruited; mean age 26.9±6.4, 29.5% male) from the study published in Kuna et al., (20) and Goldschmied et al. (13). We begin by first examining and comparing EEG signatures among unrelated individuals and within MZ and DZ twin pairs to explore their heterogeneity. We then compare MZ and DZ resemblance at progressively broader levels: first for individual traits to identify specific characteristics showing evidence of genetic influence, then jointly within related trait groups to assess evidence across shared aspects of sleep physiology, and finally across all traits to evaluate genetic influence on the combined sleep EEG phenotype. Overall, this approach aims to clarify whether genetic influence is concentrated in particular features or distributed more broadly across transient sleep oscillations, extending beyond traditionally studied fast-spindle characteristics to other frequency bands and the temporal organization of fast spindles.

## METHODS

### Participants

The study was originally approved by University of Pennsylvania Institutional Review Boards. Polysomnography data was acquired from sites subcontracted by the University of Chicago and Virginia Commonwealth University (20). Written informed consent was obtained from each participant. 53 monozygotic (MZ) and 35 dizygotic (DZ) same-sex twin pairs between ages 18 and 55 (mean age 26.9±6.4, 29.5% male) were recruited. DNA-based polymerase chain reaction analysis of peripheral blood was used to determine twin zygosity via previously described methods. All participants were screened for obstructive sleep apnea, periodic limb movements of sleep, abnormal sleep schedules, and psychiatric and neurological disease. A blood sample was also taken to exclude participants with abnormal panels. Please see details of the dataset from Kuna et al., (20) and Goldschmied et al. (13).

### EEG Recording and Preprocessing

#### Channel Selection

EEG was recorded from central electrodes referenced to the contralateral mastoids (C3–M2 and C4–M1) from multi-site clinical recordings (for details see Kuna et al., (20) and Goldschmied et al. (13)). Both the C3 and C4 channels for all twins were manually inspected. Recordings with obvious line noise or other obstructive artifacts were noted. 4 twin pairs were excluded due to one individual having errors in both channels. For analysis, C3 was chosen for all participants by default but switched to C4 when a participant’s C3 recording had a problem. If one twin was switched to C4, the other was switched as well so both twins in a pair always use the same channel. Pairs that couldn’t be matched were excluded. 5 pairs total were excluded due to channel conflicts. The final sample included 79 total pairs, comprising 47 MZ and 32 DZ twin pairs.

#### Sleep Staging, Signal Processing, and Artifact Rejection

Given variable sampling rates between recordings, all subjects were resampled to 100Hz. Staging was done manually in the original study according to AASM guidelines. Artifacts were detected automatically and removed using a high-frequency criterion 3.5 (35 Hz passband), broadband criterion 3.5 (2 Hz passband), and a 2 second smoothing window, via methods previously described in the DYNAM-O toolbox (8, 21). Only segments of sleep marked NREM 2 or NREM 3 were used for the current study.

### Feature Extraction

#### Time-Frequency Peak Detection and Parameterization

For each individual recording, we used the DYNAM-O toolbox (21) to detect and categorize all spindle-like events in the 2-16Hz frequency range. Definitionally, any spindle-like transient oscillation will appear in the time-frequency domain as a peak in the spectrogram. The DYNAM-O toolbox identifies statistically salient time-frequency peaks in the spectrogram (22) across all frequencies, and characterizes features for each extracted peak, including mean frequency, prominence, volume, boundaries, time, duration, and bandwidth. Since these events are detected agnostic to frequency, we are able to look at events that fall outside the typically analyzed spindle ranges.

The detected peaks are used to construct slow-oscillation power histograms (SOPHs) of event density (peaks per bin per minute spent at that power level) as a function of frequency and slow-oscillation (SO) power, a continuous measure of sleep depth (8, 21, 23). Events with similar properties form clusters of high density, or “modes,” on these SOPHs. Each mode represents a distinct class of spindle-like event activity, which we differentiate by relative frequency band, and the number and location of these modes are highly individualized. To characterize each mode, we fit the SOPH with a parametric model built from Gaussian basis functions. Basis functions are initialized from peaks detected on the SOPH and added one at a time, and each is kept only if it improves the model fit (8). Each fitted Gaussian defines an ellipse for each mode, which are then assigned to the previously described frequency bands according to their central frequency. For each mode, several characteristics are then extracted, including density, central frequency, SO-power, SO-phase, and mean event duration and prominence. See details in the following sections and Supplementary Table 1.

#### Event Features by Frequency Band

Previous work has shown that spindle activity as well as the activity of other spindle-like events is highly individualized (8, 24). As a result, using generalized bounds for frequency ranges may obscure meaningful differences between subjects. To address this, we identify subject-specific event characteristics. Within the power and phase histograms, clusters of similar events in frequency and sleep depth will form salient increased density modes within the histogram structure. Using the DYNAM-O toolbox, we extracted individualized features of the identified mode for each subject. Based on the center frequency of each detected mode, they were characterized into the following classes: sigma-high (11-16Hz), sigma-low (9-11Hz), alpha-theta (5-9Hz), and delta (2-5Hz). For each detected mode, the following features were extracted from the parametrized centroid: density, frequency, SO-power (sleep depth), preferred coupling in SO-phase. The peaks that fell within each detected mode’s boundaries were used to compute event-level mean features of duration, and prominence (spectral power above baseline, an analog to amplitude). If a subject has more than one mode in a given frequency band, the mode with the largest density is chosen. In instances where a mode was not identified within a given band for either twin, features for both twins were derived from individual events within the canonical band. This treats the lack of a mode as a usable observation rather than missing data.

Preferred SO phase is a circular variable. For each band, each subject’s preferred phase was centered on the circular mean across all twins and wrapped to (−π, π], so that phase differences between subjects are measured along the shortest arc and a linear ICC can be applied. The centered phases were concentrated about the mean, such that wrap-around had negligible influence on the ICC.

#### Spectral Power Features

Spectral power was computed in each band relative to mean power in the 4.5–35 Hz range in artifact-free N2 and N3 sleep periods. This accounted for non-physiological variation in baseline powers across participants likely due to recording settings. All DYNAM-O spectral-based features are inherently relative, so they were unaffected by baseline variability.

#### Timing Pattern Features

Our previous work shows that spindle timing pattern is the strongest statistical predictor of moment-to-moment spindle occurrence, surpassing the contributions of SO phase and sleep stage, which have traditionally been emphasized as key regulators of spindle activity (10). These timing patterns are highly heterogeneous across individuals yet remain relatively stable within individuals across nights. They also show age-related differences, including greater between-individual variability and shorter refractory periods in older adults. Thus, these observations position spindle timing pattern as a fingerprint-like feature of sleep neurophysiology, but whether these individual differences reflect genetic influence remains unclear. Examining their heritability may therefore reveal whether inherited contributions to spindle activity extend beyond static spindle characteristics to the dynamic, intrinsic properties of spindle production.

In this study, we estimate timing patterns through a point process framework using generalized linear models (10, 25, 26). We model the probability of seeing transient events in sigma high (fast spindles) as a function of sleep stage and past history of these events. After model fitting, the model estimates of history components can be visualized as a history modulation plot, or timing pattern, showing how past spindles modulate current spindle activity. We then quantify key features of the timing pattern, including refractory period (the duration after a spindle during which activity is significantly suppressed), peak width (the duration after the refractory period during which activity is enhanced), peak time (the lag when the maximal excitation occurs), and peak height (the degree to which the excitation is maximally enhanced). See details of model form, model fitting, choices of parameters, and implementation details in Chen et al (10). Detailed definition for all 32 traits is provided in Supplementary Table 1.

### Data Transformation

Box-Cox transformation (27) was applied to all data, fitted per trait and applied only where its lambda CI excluded 1, with dB and phase traits skipped. The SO phase was centered on its circular mean.

### Statistical Analysis

#### Intraclass Correlations

Twin resemblance for each trait was measured with the one-way random-effects intraclass correlation, ICC(1,1). It was computed separately for MZ and DZ pairs. This form treats the two co-twins as interchangeable, so the result does not depend on which twin is listed first. The ICC was calculated from the between-pair and within-pair mean squares of a one-way ANOVA. Its 95% confidence interval was taken from the F distribution with n − 1 and n degrees of freedom, where n is the number of pairs. For each trait, only pairs with data for both twins were included. To characterize the correlation structure among traits, we computed the correlation matrix of all traits across subjects.

#### Comparing MZ and DZ Correlations

A greater resemblance in MZ than in DZ twins is the basic requirement for a genetic interpretation of twin data. Attributing this difference to genetic influence assumes that MZ and DZ pairs share environments to the same degree (the equal-environments assumption). For each trait, the difference between the MZ and DZ ICCs was expressed as a Fisher z statistic:

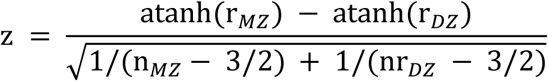

where n is the number of complete pairs and 1/(n − 3/2) is the large-sample variance of the transformed intraclass correlation. This puts every trait on a common, variance-stabilized scale.

#### Permutation Testing and Multiple Comparisons

The Fisher z statistic is referred to the standard normal distribution only in large samples. In our data its permutation null distribution had wider tails, so p-values from the analytic test would be anticonservative. We therefore assessed significance by permutation. The MZ and DZ labels were shuffled across pairs 100,000 times, keeping each pair together and the numbers of MZ and DZ pairs fixed, and the Fisher z of every trait was recomputed under the same shuffled labels. For each trait, the one-sided empirical p-value was (1 + b)/(1 + B), where b is the number of the B = 100,000 permuted values at or above the observed one.

The empirical p-values were adjusted for multiple comparisons across the 32 traits with the Benjamini-Hochberg false discovery rate (FDR) procedure (28). Because of the small sample size, these analyses were treated as exploratory: adjusted p-values are reported for all traits to rank the effects, and no traits were selected or dropped on the basis of a significance threshold.

#### Group Tests Across Trait Categories

To test whether MZ twins resembled each other more than DZ twins across a group of traits, rather than one trait at a time, we used three group statistics. The mean Fisher z across the traits in the group is sensitive to small effects spread over many traits. The maximum Fisher z is sensitive to a strong effect in a few traits, and because its null is the distribution of the largest value across the group, it controls the familywise error rate. Both were evaluated on the same 100,000 label shuffles as the single-trait tests, so the correlations among traits are preserved in their null distributions. The third statistic compared whole trait profiles: each trait was z-scored across all twins, the co-twin distance of each pair was computed as the root-mean-square difference across the group’s traits, and the statistic was the mean DZ distance minus the mean MZ distance. Pairs missing any trait in the group were excluded from that group’s distance test. Its null distribution was built from 100,000 separate shuffles of the pair labels under the same scheme.

The traits were organized into three blocks:

1. Mode features, the six event features (density, frequency, SO power, phase preference, duration, and prominence) in each of the four frequency bands
2. The four parameters of the fast-spindle (sigma-high) timing pattern (refractory period, peak time, peak height, and peak width)
3. Relative spectral power in each band (mean power in the band relative to 4.5–35 Hz, in dB).

In the mode block, group tests were run for each band (its six features), for each feature across the four bands, and for the whole block. The timing and spectral blocks, which have a single row and a single column respectively, were tested as whole blocks. A global test was run on all 32 traits. Group results are reported as empirical p-values, and as permutation z-scores.

#### Heritability Model Estimates

While direct estimates of heritability are limited in power for low-N studies, we computed supplementary Falconer’s estimates (29) in order to align with previous literature, and to understand what proportion of the variance of each trait can be attributed to additive genetics (h²), shared environments (c²), or unique environments (e²). The ACE model was forced (30) (as opposed to having an ADE option and then performing a likelihood ratio test) due to the small sample size. Estimates were not constrained to the 0-1 range and are reported in Supplementary materials exactly as they were calculated.

## RESULTS

### Individualized sleep EEG signatures show markedly reduced heterogeneity in twins

Given previous findings of strong intersubject heterogeneity in high-dimensional sleep EEG signatures, even within tightly triaged cohorts (8), it is important to determine the degree to which this variability appears to be reduced within twin pairs. In Figure 1, we demonstrate the variety of transient oscillatory brain signatures seen in an illustrative subset of the subjects from the twin study. Subjects are broken into unrelated, MZ, and DZ categories. For each subject, we show the slow-oscillation power histogram (SOPH).

**Figure 1.**
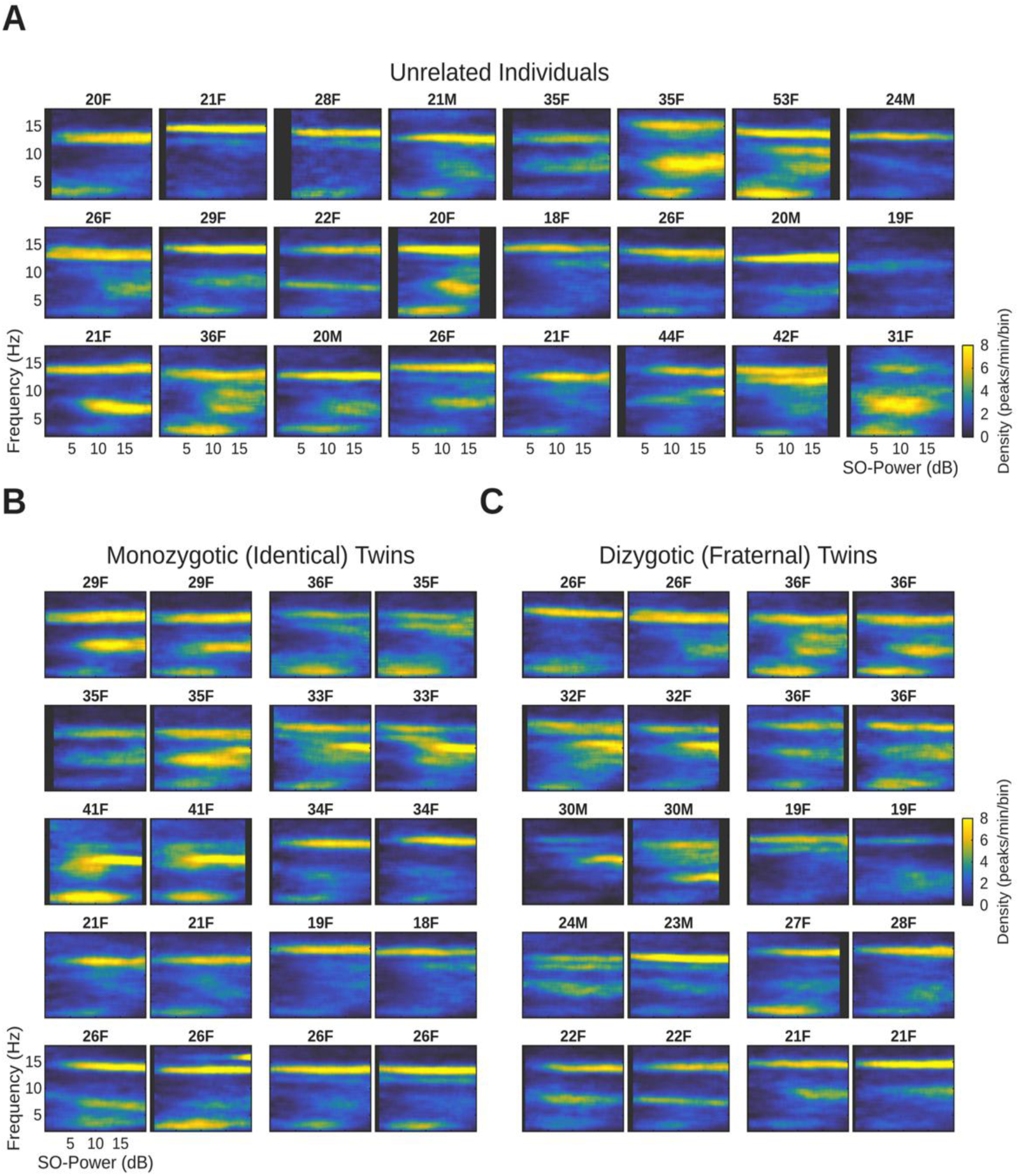
Monozygotic twins show greater similarity in slow oscillation power histograms (SOPH) than dizygotic twins and unrelated individuals. In each SOPH, color represents the density (event/min/bin) of spindle-like events as a function of frequency (y-axis) and sleep depth, as indexed by SO-power (x-axis). In Panel A, a random selection of 32 non-related participants from the dataset are shown. The bottom panels show monozygotic (B, left) and dizygotic (C, right) twin pairs, labeled with age (in years) and sex (F: female; M: male).

Each SOPH displays the rate of detected transient oscillations across frequency and slow-oscillation (SO) power (left to right indicates light to deep NREM), allowing transient events to be examined across continuous sleep depth. In Figure 1A, the unrelated participants exhibit substantial variation in the central frequency, SO-power location, and relative prominence of event concentrations, including activity outside the traditional sigma range associated with sleep spindles. Moreover, the number of modes differs between different subjects. This intersubject heterogeneity is consistent with previous studies (8). In contrast, twin pairs show markedly similar distributions, with particularly close correspondence apparent among MZ pairs (Figure 1B). The number of modes appears to be generally conserved within twin pairs. These examples suggest that twins show strong similarity across a set of high-dimensional features, which taken together, may prove more informative than any single feature in isolation.

### Trait-level analyses reveal genetic influence beyond sigma-high range

Having observed reduced heterogeneity within twin pairs, we next ask which individual characteristics show evidence of genetic influence and whether that evidence extends beyond sigma-high activity, which corresponds to fast spindles. To address these questions, we compare trait-specific intraclass correlations (ICCs) between MZ and DZ pairs using permutation tests, with false discovery rate correction across all 32 traits. The ICC quantifies within-pair resemblance relative to between-pair variation, with higher values indicating greater similarity between co-twins. Five of the 24 mode characteristics show significantly greater MZ than DZ resemblance after correction (Table 1).

**Table 1.** reports trait-level comparisons of MZ and DZ intraclass correlations for each frequency band, with adjustment for multiple comparisons. ICC, intraclass correlation. CI, confidence interval. Z denotes the Fisher’s z-test statistic. P-values test whether the MZ ICC exceeds the DZ ICC using 100,000 permutations of zygosity labels across twin pairs, and adjusted P-values are computed using the Benjamini–Hochberg procedure across all 32 traits. Throughout this work, significance levels are indicated by dark green (P < 0.001), medium green (P < 0.01), light green (P < 0.05), and yellow (P < 0.10) shading, respectively.

| Feature | ICC<sub>MZ</sub> | 95% CI | ICC<sub>DZ</sub> | 95% CI | Z | P-value | Adjusted P-value |
|---|---|---|---|---|---|---|---|
| Sigma High |  |  |  |  |  |  |  |
| Density | 0.6778 | [0.4888, 0.8063] | 0.3050 | [-0.0396, 0.5861] | 2.1790 | 0.0281 | 0.1123 |
| Frequency | 0.6233 | [0.4135, 0.7706] | 0.7110 | [0.4889, 0.8471] | -0.6786 | 0.7140 | 0.7878 |
| SO power | 0.5482 | [0.3142, 0.7198] | -0.1041 | [-0.4294, 0.2469] | 3.0779 | 0.0061 | 0.0414 |
| Phase Pref | 0.0743 | [-0.2128, 0.3504] | 0.1564 | [-0.1944, 0.4734] | -0.3561 | 0.6206 | 0.7215 |
| Duration | 0.8218 | [0.7023, 0.8964] | 0.3052 | [-0.0394, 0.5863] | 3.6194 | 0.0016 | 0.0414 |
| Prominence | 0.8147 | [0.6913, 0.8921] | 0.3818 | [0.0475, 0.6404] | 3.1565 | 0.0052 | 0.0414 |
| Ref period | 0.6783 | [0.4871, 0.8078] | 0.1629 | [-0.1880, 0.4785] | 2.8146 | 0.0063 | 0.0414 |
| Peak time | 0.4656 | [0.2075, 0.6635] | 0.2406 | [-0.1088, 0.5385] | 1.1017 | 0.1513 | 0.3201 |
| Peak height | 0.6717 | [0.4777, 0.8035] | 0.2459 | [-0.1033, 0.5425] | 2.3940 | 0.0160 | 0.0732 |
| Peak width | 0.3126 | [0.0296, 0.5498] | -0.0592 | [-0.3918, 0.2890] | 1.6277 | 0.0504 | 0.1791 |
| Mean power | 0.7067 | [0.5299, 0.8249] | 0.4728 | [0.1577, 0.7015] | 1.5678 | 0.0748 | 0.2176 |
| Sigma Low |  |  |  |  |  |  |  |
| Density | 0.7443 | [0.5847, 0.8487] | 0.6655 | [0.4203, 0.8208] | 0.6729 | 0.2858 | 0.5380 |
| Frequency | 0.5343 | [0.2964, 0.7102] | 0.4428 | [0.1205, 0.6817] | 0.5145 | 0.4158 | 0.5785 |
| SO power | 0.5569 | [0.3255, 0.7258] | -0.0676 | [-0.3989, 0.2812] | 2.9740 | 0.0065 | 0.0414 |
| Phase Pref | 0.3642 | [0.0909, 0.5870] | 0.5382 | [0.2421, 0.7435] | -0.9398 | 0.8306 | 0.8306 |
| Duration | 0.8523 | [0.7505, 0.9148] | 0.7098 | [0.4870, 0.8464] | 1.6149 | 0.0630 | 0.2017 |
| Prominence | 0.4224 | [0.1587, 0.6304] | 0.4085 | [0.0790, 0.6587] | 0.0717 | 0.4687 | 0.5957 |
| Mean power | 0.7716 | [0.6254, 0.8657] | 0.7532 | [0.5550, 0.8709] | 0.1884 | 0.4388 | 0.5850 |
| Alpha-Theta |  |  |  |  |  |  |  |
| Density | 0.7529 | [0.5974, 0.8541] | 0.6289 | [0.3669, 0.7991] | 1.0262 | 0.1601 | 0.3201 |
| Frequency | 0.4802 | [0.2286, 0.6722] | 0.6315 | [0.3708, 0.8007] | -0.9432 | 0.7630 | 0.8138 |
| SO power | 0.5706 | [0.3434, 0.7352] | 0.0169 | [-0.3254, 0.3571] | 2.6987 | 0.0091 | 0.0483 |
| Phase Pref | 0.3527 | [0.0778, 0.5784] | 0.2283 | [-0.1217, 0.5292] | 0.5819 | 0.3066 | 0.5451 |
| Duration | 0.7906 | [0.6541, 0.8774] | 0.6067 | [0.3355, 0.7858] | 1.5779 | 0.1054 | 0.2530 |
| Prominence | 0.5219 | [0.2807, 0.7016] | 0.6706 | [0.4278, 0.8238] | -0.9948 | 0.8294 | 0.8306 |
| Mean power | 0.7160 | [0.5433, 0.8308] | 0.6667 | [0.4220, 0.8215] | 0.4041 | 0.3469 | 0.5690 |
| Delta |  |  |  |  |  |  |  |
| Density | 0.5760 | [0.3504, 0.7388] | 0.5152 | [0.2119, 0.7289] | 0.3704 | 0.3734 | 0.5690 |
| Frequency | 0.6186 | [0.4072, 0.7675] | 0.2941 | [-0.0516, 0.5782] | 1.7935 | 0.0873 | 0.2329 |
| SO power | 0.3378 | [0.0610, 0.5670] | -0.0212 | [-0.3591, 0.3234] | 1.5932 | 0.1107 | 0.2530 |
| Phase Pref | 0.2585 | [-0.0261, 0.5049] | 0.3571 | [0.0189, 0.6232] | -0.4662 | 0.6313 | 0.7215 |
| Duration | 0.3672 | [0.0943, 0.5893] | 0.3588 | [0.0209, 0.6244] | 0.0410 | 0.4840 | 0.5957 |
| Prominence | 0.3819 | [0.1113, 0.6004] | 0.3098 | [-0.0343, 0.5896] | 0.3502 | 0.3669 | 0.5690 |
| Mean power | 0.7157 | [0.5428, 0.8306] | 0.6750 | [0.4343, 0.8264] | 0.3368 | 0.3927 | 0.5713 |

The clearest evidence occurs in sigma-high mode features. Mode duration and prominence show substantially greater resemblance among MZ twins, with MZ versus DZ ICCs of 0.82 versus 0.31 for duration and 0.81 versus 0.38 for prominence. The SO-power of sigma-high modes also shows greater MZ resemblance, with ICCs of 0.55 versus −0.10. All three characteristics remain significant after correction (adjusted P-value = 0.041 for each; Table 1). These results are consistent with earlier twin and family studies reporting heritability of spindle characteristics in the sigma range (9, 13, 14), and further support studies showing that spindle heritability estimates are robust to the choice of detector (13).

Heritability evidence also extends to several characteristics of lower-frequency activity. Specifically, SO-power shows greater MZ than DZ resemblance in both sigma-low activity (ICC = 0.56 versus −0.07; adjusted P-value = 0.041) and alpha-theta activity (ICC = 0.57 versus 0.02; adjusted P-value = 0.048). Thus, although the significance of mode characteristics is more prevalent in the sigma-high band, SO-power shows evidence of genetic influence across three frequency bands, extending beyond the spindle range.

No mode density, frequency, or SO-phase preference trait shows significantly greater MZ than DZ resemblance after correction. Likewise, none of the relative spectral-power measures survives correction (Table 1).

### Spindle coupling with the slow oscillation phase does not show a measurable genetic influence

Twin similarity in the preferred SO phase of events has not, to our knowledge, been examined before. Unlike duration, prominence and SO power, preferred phase showed no evidence that MZ twins were more similar than DZ twins in any band (Fisher z -0.94 to 0.58; P-value = 0.31 to 0.83; Table 1). Co-twin resemblance in phase nonetheless differed by band. In sigma low, preferred phase was moderately correlated within pairs, and if anything more so in DZ pairs (rMZ = 0.36, rDZ = 0.54). In sigma high, it was barely correlated within either group (r_MZ_ = 0.07, r_DZ_ = 0.16). Delta (r_MZ_ = 0.26, r_DZ_ = 0.36) and alpha/theta (r_MZ_ = 0.35, r_DZ_ = 0.23) fell in between. These results provide no evidence of genetic influence on preferred SO phase, although phase in sigma low appears to be familial, possibly reflecting shared environmental or measurement factors. The low within-pair correlations in sigma high suggest that preferred phase in that band may be too noisy to estimate precisely from a single night.

### Fast spindle timing patterns show evidence of genetic influence

We next examine whether genetic influences extend to the timing patterns of sigma-high events (traditional fast spindles), an important but often overlooked aspect of spindle activity. Our previous work illustrates that timing patterns play a dominant role in spindle production, yet their heritability has not been characterized. We estimate fast spindle timing patterns for each individual and summarize them using four interpretable parameters: refractory period, peak time, peak height, and peak width. The ICC and its 95% confidence intervals are shown in Table 1.

Refractory period, which describes the interval of suppressed event occurrence following a preceding spindle, shows substantially greater resemblance among MZ than DZ twins. The MZ ICC is 0.68 (95% CI: [0.4871, 0.8078]), compared with 0.16 (95% CI: [-0.1880, 0.4785]) in DZ twins, and this difference remains significant after permutation testing and false discovery rate correction across all 32 traits (adjusted P-value = 0.041). Timing pattern peak height, which measures the maximum enhancement of event occurrence following the refractory period, also shows greater MZ resemblance (ICC = 0.67 versus 0.25). However, this difference does not survive correction (P-value = 0.016; adjusted P-value = 0.073). Neither peak time nor peak width survives correction (adjusted P-value = 0.32 and 0.18, respectively).

### MZ twins are significantly closer than DZ twins in high-dimensional feature space

Having examined individual traits separately and related traits jointly within groups, we now investigate whether MZ co-twins show greater overall resemblance than DZ co-twins when all 32 traits are considered together. To quantify overall resemblance, we calculate Euclidean pairwise distances using all 32 standardized traits as dimensions for all subjects in the population. Figure 2 shows the distribution of pairwise distances between all MZ co-twins, all DZ co-twins, and all unrelated pairs in the population. The mean distance is smallest for MZ pairs (0.86), followed by DZ pairs (1.14) and unrelated pairings (1.38), with smaller distances indicating greater similarity. MZ co-twins are significantly closer than DZ co-twins (DZ–MZ difference = 0.28; P-value = 0.00002, permutation test; Table 2). Both twin groups also have smaller mean distances than the unrelated reference pairings. A detailed table of key statistics for Figure 2 is in provided in Supplementary Table 3.

**Figure 2.**
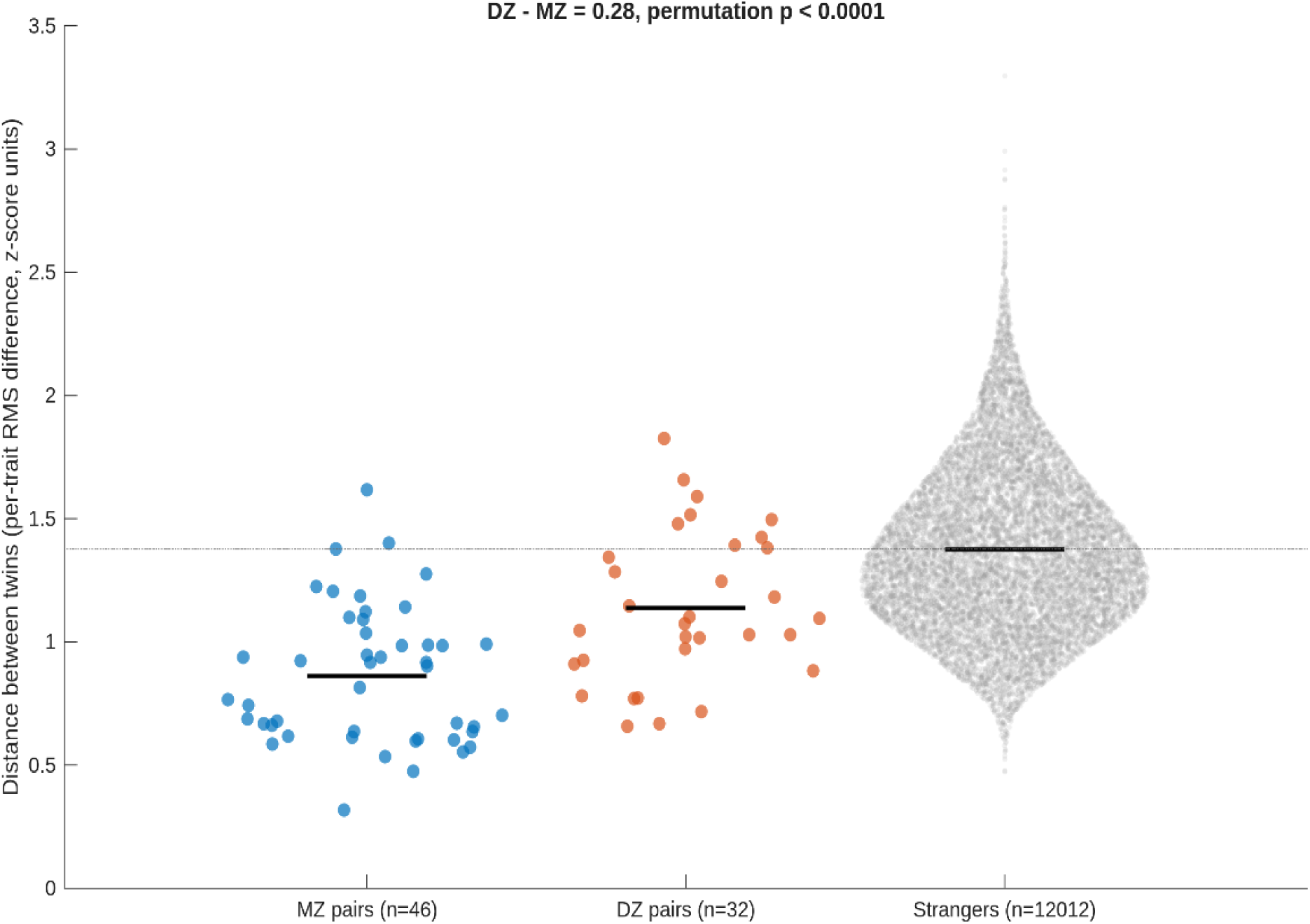
MZ co-twins show significantly greater multivariate similarity than DZ co-twins and unrelated pairs. Each point represents the distance between two individuals across all 32 sleep EEG traits, shown for MZ twin pairs (blue, n = 46), DZ twin pairs (orange, n = 32) and all unrelated pairings (“strangers”, gray, n = 12,012). Distance is the root-mean-square difference per trait in Z-score units. Black bars mark group means, and the dotted horizontal line marks the stranger mean as a reference line. MZ co-twins were closer than DZ co-twins, and both are closer than unrelated pairs.

**Table 2.** P-values for individual-trait and joint comparisons of MZ and DZ resemblance across trait groups and all 32 traits.

| Group | Density | Frequency | SO power | Phase Pref | Duration | Prominence | Max Z | Mean Z | Pair Dist |
|---|---|---|---|---|---|---|---|---|---|
| Sigma high (11-16 Hz) | 0.0281 | 0.714 | **0.00612** | 0.621 | **0.0016** | **0.00517** | **0.00541** | **0.00196** | **0.00445** |
| Sigma low (9-11 Hz) | 0.286 | 0.416 | **0.00647** | 0.831 | 0.063 | 0.469 | 0.15 | 0.106 | **0.00372** |
| Alpha/theta (5-9 Hz) | 0.16 | 0.763 | **0.00906** | 0.307 | 0.105 | 0.829 | 0.0464 | 0.1 | 0.0211 |
| Delta (2-5 Hz) | 0.373 | 0.0873 | 0.111 | 0.631 | 0.484 | 0.367 | 0.308 | 0.121 | **0.00079** |
| Across bands: Max Z | 0.101 | 0.398 | 0.0129 | 0.743 | **0.00304** | 0.0144 | – | – | – |
| Across bands: Mean Z | 0.0328 | 0.406 | **0.00754** | 0.68 | **0.00055** | 0.152 | – | – | – |
| Across bands: Pair dist | **0.00274** | 0.562 | **0.00174** | 0.762 | **4e-05** | 0.0224 | – | – | – |
| **Block aggregate (24 traits)** | – | – | – | – | – | – | 0.0973 | **0.00145** | **0.00016** |
| **Timing pattern (sigma high)** |  |  |  |  |  |  |  |  |  |
| **Group** | **Ref period** | **Peak time** | **Peak height** | **Peak width** | **Max Z** | **Mean Z** | **Pair Dist** |  |  |
| Sigma high (11-16 Hz) | **0.00632** | 0.151 | 0.016 | 0.0504 | – | – | – |  |  |
| **Block aggregate (4 traits)** | – | – | – | – | 0.0157 | **0.00016** | 0.0191 |  |  |
| **Spectral power** |  |  |  |  |  |  |  |  |  |
| **Group** | **Mean Spectral Power** | **Max Z** | **Mean Z** | **Pair Dist** |  |  |  |  |  |
| Sigma high (11-16 Hz) | 0.0748 | – | – | – |  |  |  |  |  |
| Sigma low (9-11 Hz) | 0.439 | – | – | – |  |  |  |  |  |
| Alpha/theta (5-9 Hz) | 0.347 | – | – | – |  |  |  |  |  |
| Delta (2-5 Hz) | 0.393 | – | – | – |  |  |  |  |  |
| **Block aggregate (4 traits)** | – | 0.281 | 0.154 | 0.137 |  |  |  |  |  |
| **Aggregates by block and overall** |  |  |  |  |  |  |  |  |  |
| **Group** | **Max Z** | **Mean Z** | **Pair Dist** |  |  |  |  |  |  |
| Mode parameters (24 traits) | 0.0973 | **0.00145** | **0.00016** |  |  |  |  |  |  |
| Timing pattern (4 traits) | 0.0157 | **0.00016** | 0.0191 |  |  |  |  |  |  |
| Spectral power (4 traits) | 0.281 | 0.154 | 0.137 |  |  |  |  |  |  |
| **GRAND AGGREGATE (32 traits)** | 0.1 | **9e-05** | **2e-05** |  |  |  |  |  |  |

Individual cells report one-sided permutation P-values testing whether the MZ ICC exceeds the DZ ICC, based on 100,000 permutations of zygosity labels across twin pairs. Max Z, Mean Z, and Pair Dist (pairwise distance) denote group tests based on the maximum Fisher z-statistic, mean Fisher z-statistic, and co-twin distance, respectively, across the traits in each row, each column (“Across bands” rows), or the entire block. Shading indicates significance levels and “–” denotes not applicable.

Together, the greater overall resemblance among MZ than DZ twins reinforces the evidence from individual traits and related trait groups, supporting genetic influence across the combined sleep EEG phenotype. These findings are consistent with genetic influence across a broad set of transient sleep oscillations and their distinct neural signatures, extending beyond the spindle characteristics examined in previous heritability studies (9, 13, 14).

### Joint tests reveal stronger MZ–DZ differences than individual traits

To assess evidence across groups of related traits, we tested each trait block (mode parameters, timing pattern and spectral power) and the full set of 32 traits using three complementary statistics. The maximum-Z test evaluates the strongest single-trait Fisher z statistic within a group, the mean-Z test averages the Fisher z statistic across the group’s traits, and co-twin distance compares the resemblance of MZ and DZ pairs across the group’s combined standardized trait profile. Each statistic was evaluated against its permutation null distribution. Because co-twin distance is computed on a scale pooled across zygosity, it reflects both within-pair resemblance and any difference in trait variance between the MZ and DZ groups.

Results are summarized in Table 2. The first trait category is the SOPH mode parameters, organized with frequency bands as rows and characteristics as columns. Marginal summaries report the joint tests for the six characteristics within each frequency band (row-wise) and for each characteristic across frequency bands (column-wise). Combining the six mode characteristics within each band, MZ twins showed smaller co-twin distances than DZ twins in all four bands (sigma high P-value = 0.0045, sigma low P-value = 0.0037, alpha/theta P-value = 0.021, delta P-value = 0.00079). Only sigma high also reached significance on the mean-Z test (P-value = 0.0020; other bands P-value = 0.10–0.12).

Event density illustrates the distinction between individual and joint analyses. MZ correlations exceeded DZ correlations in all four bands (rMZ − rDZ = 0.06–0.37), but no single band survived FDR correction (smallest adjusted P-value = 0.11, sigma high), and the maximum-Z test across bands did not reach nominal significance (P-value = 0.10). Tested jointly, however, density showed a significant MZ–DZ difference in both the mean-Z test (P-value = 0.033) and co-twin distance (P-value = 0.0027). The combined result therefore reflects a consistent small effect spread across bands rather than a strong effect in one band.

Joint analyses also identified differences when the same characteristic was considered across frequency bands. Although only sigma-high duration survived individual-trait correction, duration combined across all four bands showed greater MZ resemblance under the maximum-Z (P-value = 0.0030), mean-Z (P-value = 0.00055) and co-twin distance tests (P-value = 0.00004). SO power also showed evidence across bands under all three tests (maximum-Z P-value = 0.013, mean-Z P-value = 0.0075, co-twin distance P-value = 0.0017). Across all 24 mode traits, MZ twins showed greater resemblance under both the mean-Z test (P-value = 0.0015) and the co-twin distance test (P-value = 0.00016), whereas the maximum-Z test did not reach significance (P-value = 0.097).

The next trait category comprises the four parameters of the fast-spindle (sigma-high) timing pattern. Considered together, these parameters showed greater MZ than DZ resemblance under the maximum-Z (P-value = 0.016), mean-Z (P-value = 0.00016) and co-twin distance tests (P-value = 0.019). These findings extend the evidence of genetic influence from individual timing parameters (e.g., refractory period, adjusted P-value = 0.041) to the timing pattern assessed jointly. We also tested the four spectral power measures jointly. None of the three tests showed significant evidence of greater MZ than DZ resemblance (maximum-Z P-value = 0.28, mean-Z P-value = 0.15, co-twin distance P-value = 0.14).

Finally, in the omnibus test across all 32 traits, both the mean-Z test (P-value = 0.00009) and the co-twin distance test (P-value = 0.00002) showed significantly greater MZ than DZ resemblance, whereas the maximum-Z test did not reach significance (P-value = 0.10). Because the mean-Z test aggregates evidence across traits while the maximum-Z test evaluates the strongest individual effect, this pattern is consistent with genetic influence distributed across many traits rather than concentrated in a single dominant trait. The same comparisons expressed as permutation z-scores, which standardize each observed statistic against its permutation null distribution, show a similar pattern (Supplementary Table 2). Thus, although few individual traits survive correction, joint analyses of related trait groups and of the full set of 32 traits provide strong evidence of greater MZ than DZ resemblance, consistent with genetic influence on the overall oscillatory phenotype.

## DISCUSSION

### Genetic influence is distributed across traits and most detectable in the combined phenotype

This study provides additional evidence of genetic influence on previously unexplored features of transient sleep oscillations, including their distribution across frequency and sleep depth and the temporal organization of fast spindles. The evidence strengthens as analyses progress from individual traits to related trait groups and ultimately to the combined sleep EEG phenotype. At the individual-trait level, greater MZ than DZ resemblance extends beyond sigma-high activity to features of lower-frequency oscillations, while analysis of fast-spindle timing identifies genetic influence on the refractory period. Joint analyses of several related trait groups provide stronger evidence, revealing broader patterns of resemblance that are less apparent when traits are examined separately. The strongest evidence emerges when all 32 traits are considered together: the mean-Z and co-twin-distance tests yield P-values one to two orders of magnitude smaller than the smallest uncorrected single-trait P-value. Together, these findings suggest that genetic influence is distributed across multiple oscillatory characteristics, extending beyond traditionally studied fast-spindle features, and is most clearly detected when the broader sleep EEG phenotype is considered as a whole.

### Genetic influence on transient oscillations extends beyond the fast-spindle band

Among the SOPH mode parameters, sigma-high duration and prominence show among the strongest evidence of genetic influence (Table 1). These findings corroborate previous studies reporting heritable variation in conventional spindle characteristics. Goldschmied et al. (13) found robust genetic contributions to spindle characteristics across multiple detection algorithms, while demonstrating that the evidence varied by characteristic and detector. Moreover, our absence of a significant MZ–DZ difference in mean frequency is consistent with their less uniform evidence for spindle frequency, for which significant additive genetic variance was detected with only two of eight algorithms (13). Evidence also extends beyond sigma-high activity: SO-power shows greater MZ resemblance in sigma-high, sigma-low, and alpha-theta bands. Building on the characterization of oscillatory activity across continuous sleep depth, these findings suggest that genetic influences extend to the sleep-depth conditions under which transient events concentrate.

### Spindle timing is a newly identified genetically influenced trait

A further contribution is the evidence of genetic influence on intrinsic spindle timing patterns, as captured by the dependence of event occurrence on spindle history. Our previous work demonstrated that these timing patterns are highly individualized, relatively stable across nights, and provide substantial predictive information about subsequent spindle occurrence (10). Here, the refractory period shows significantly greater MZ than DZ resemblance after correction, suggesting a genetic contribution to individual differences in the interval of suppressed event occurrence following a spindle (Table 1). Although the other timing parameters do not individually survive correction for multiple comparisons, joint analyses provide evidence of genetic influence on the timing parameters considered collectively (Table 2). The potential functional importance of this result is supported by a previous study showing that memory cues delivered just after the spindle refractory period produced better memory outcomes than cues delivered within it (31). Therefore, these observations motivate future investigation of whether genetically influenced variation in spindle timing contributes to individual differences in sleep-dependent memory processing.

### Limited power constrains null findings and ACE estimates

Several characteristics, including preferred SO phase, mean frequency and relative spectral power, showed no significant MZ–DZ differences either individually or jointly, and density was significant only when tested jointly across bands. These findings should be interpreted cautiously, as a lack of statistical significance does not establish the absence of genetic influence. Although this is among the largest twin sleep EEG cohorts studied to date, detecting MZ–DZ differences requires sufficiently precise ICC estimates in both groups, and correction for multiple comparisons further limits power to detect modest individual-trait effects.

As an exploratory analysis to assess how co-twin resemblance may reflect different sources of variation, we calculated Falconer estimates of additive genetic, shared environmental, and unique environmental contributions under the ACE framework (30). In Supplementary Table 4, we see that sigma-low relative spectral power shows similarly high ICCs in MZ and DZ twins (0.77 and 0.75, respectively), yielding estimates of 0.04 for the additive genetic component and 0.73 for the shared environmental component. Under the ACE assumptions, this decomposition attributes the strong co-twin resemblance primarily to shared environmental influences. However, more than half of the traits (17 of 32) yield at least one estimate outside the valid range, including additive genetic estimates above 1 (up to 1.30 for sigma-high SO power), and negative shared environmental estimates for several traits. These values cannot be interpreted literally as proportions of variance and may reflect imprecise ICC estimates in this sample, departures from the ACE assumptions, or both. The Falconer estimates therefore remain exploratory and do not support firm conclusions about the relative genetic and environmental contributions to individual traits.

### Future Work

Future study design should also account for the correlations among traits. The correlation matrix in Supplementary Figure 1 shows that several traits are strongly correlated (up to r = 0.87), making this dependence relevant to power calculations for joint analyses. A formal power analysis incorporating the observed correlation structure, a range of plausible effect sizes, and the planned multiple-comparison procedures would help determine the numbers of MZ and DZ pairs needed for individual-trait and joint tests. Such analyses would guide enrollment targets for studies aimed at detecting more modest individual-trait effects and estimating genetic and environmental contributions with greater precision.

Future work will also examine genetic influences on the spatial and temporal organization of transient sleep oscillations. Cox et al. (24) demonstrated individual differences in spindle frequency and topography, and Rusterholz et al. (32) further showed regional variation in genetic and environmental contributions to sleep neurophysiology in adolescents, providing a direct rationale for extending the present central-channel analyses to broader scalp coverage. Applying point process models of fast spindles to other frequency band events across multiple channels would further clarify whether history-dependent suppression and enhancement characterize broader transient oscillatory activity, how these temporal patterns vary across brain regions, and whether individual differences in these patterns show evidence of genetic influence.

### Conclusions

Genetic influence on sleep oscillations is not confined to a few spindle features. It is distributed across the oscillatory phenotype. Examined one trait at a time, this influence is only partly visible. Examined jointly, it emerges clearly: the combined profile of 32 traits separates MZ from DZ twins more decisively than any single trait does. Within this profile, we identify genetic influence on features that have not previously been examined, including the sleep depth at which transient events occur across frequency bands and the refractory period that governs the timing of fast spindles. These results suggest that the heritability of sleep EEG can be thought of in more holistic terms, considering the entire oscillatory system rather than isolated measures. Moreover, this study highlights that at the sample sizes achievable in twin studies, single-trait analyses can substantially understate genetic influence that joint analyses can detect. Because spindles and the oscillations that accompany them are linked to memory, aging, and neurological disease, this expanded set of genetically influenced features offers new targets for understanding how inherited differences in sleep physiology shape cognitive and clinical outcomes.

## CODE AVAILABILITY

Code for the analysis in this paper will be available upon publication.

## SUPPLEMENTARY MATERIAL

Supplementary material is provided in a separate document and will be available online.

## REFERENCES

1. A. L. Loomis, E. N. Harvey, G. Hobart, Potential rhythms of the cerebral cortex during sleep. Science 81, 597–598 (1935).

2. L. M. J. Fernandez, A. Lüthi, Sleep Spindles: Mechanisms and Functions. Physiol. Rev. 100, 805–868 (2020).

3. B. A. Mander, J. R. Winer, M. P. Walker, Sleep and human aging. Neuron 94, 19–36 (2017).

4. J. W. Antony, M. Schönauer, B. P. Staresina, S. A. Cairney, Sleep Spindles and Memory Reprocessing. Trends Neurosci. 42, 1–3 (2019).

5. D. S. Manoach, R. Stickgold, Abnormal sleep spindles, memory consolidation, and schizophrenia. Annu. Rev. Clin. Psychol. 15, 451–479 (2019).

6. C. A. Farmer, et al., Spindle activity in young children with autism, developmental delay, or typical development. Neurology 91, e112–e122 (2018).

7. M. Gorgoni, et al., Parietal Fast Sleep Spindle Density Decrease in Alzheimer’s Disease and Amnesic Mild Cognitive Impairment. Neural Plast. 2016, 8376108 (2016).

8. P. A. Stokes, et al., Transient oscillation dynamics during sleep provide a robust basis for electroencephalographic phenotyping and biomarker identification. Sleep 46, zsac223 (2023).

9. S. M. Purcell, et al., Characterizing sleep spindles in 11,630 individuals from the National Sleep Research Resource. Nat. Commun. 8, 15930 (2017).

10. S. Chen, M. He, R. E. Brown, U. T. Eden, M. J. Prerau, Individualized temporal patterns drive human sleep spindle timing. Proc. Natl. Acad. Sci. 122, e2405276121 (2025).

11. L. De Gennaro, et al., The electroencephalographic fingerprint of sleep is genetically determined: a twin study. Ann. Neurol. 64, 455–460 (2008).

12. U. Ambrosius, et al., Heritability of sleep electroencephalogram. Biol. Psychiatry 64, 344–348 (2008).

13. J. R. Goldschmied, et al., Spindles are highly heritable as identified by different spindle detectors. Sleep 44, zsaa230 (2021).

14. M. Adamczyk, L. Genzel, M. Dresler, A. Steiger, E. Friess, Automatic Sleep Spindle Detection and Genetic Influence Estimation Using Continuous Wavelet Transform. Front. Hum. Neurosci. 9, 624 (2015).

15. R. F. Helfrich, B. A. Mander, W. J. Jagust, R. T. Knight, M. P. Walker, Old Brains Come Uncoupled in Sleep: Slow Wave-Spindle Synchrony, Brain Atrophy, and Forgetting. Neuron 97, 221–230.e4 (2018).

16. M. A. Hahn, D. Heib, M. Schabus, K. Hoedlmoser, R. F. Helfrich, Slow oscillation-spindle coupling predicts enhanced memory formation from childhood to adolescence. eLife 9, e53730 (2020).

17. H. Noamany, et al., Multiple classes of spindle-like oscillations reveal stereotyped lifespan dynamics amid individual variability. Available at: 10.64898/2026.11.20.768147.

18. T. Dimitrov, M. He, R. Stickgold, M. J. Prerau, Sleep spindles comprise a subset of a broader class of electroencephalogram events. Sleep (2021). 10.1093/sleep/zsab099.

19. N. Biabani, et al., Disorder-specific alterations of transient oscillatory dynamics during sleep across cortical and subcortical networks. Sci. Rep. 16, 3630 (2026).

20. S. T. Kuna, et al., Heritability of performance deficit accumulation during acute sleep deprivation in twins. Sleep 35, 1223–1233 (2012).

21. M. He, S. R. Saremsky, H. Noamany, S. Chen, M. J. Prerau, The DYNAM-O Toolbox: Characterizing Individualized Neural Signatures in Sleep EEG. [Preprint] (2026). Available at: https://www.biorxiv.org/content/10.64898/2026.08.26.747401v1 [Accessed 2 September 2026].

22. M. J. Prerau, R. E. Brown, M. T. Bianchi, J. M. Ellenbogen, P. L. Purdon, Sleep neurophysiological dynamics through the lens of multitaper spectral analysis. Physiology 32, 60–92 (2017).

23. H. J. Kim, S. Chen, U. T. Eden, M. J. Prerau, A quantitative representation of continuous brain state during sleep in 2021 *10th International IEEE/EMBS Conference on Neural Engineering (NER)*, (2021), pp. 103–106.

24. R. Cox, A. C. Schapiro, D. S. Manoach, R. Stickgold, Individual differences in frequency and topography of slow and fast sleep spindles. Front. Hum. Neurosci. 11, 433 (2017).

25. S. Chen, S. Redline, U. T. Eden, M. J. Prerau, Dynamic Models of Obstructive Sleep Apnea Provide Robust Prediction of Respiratory Event Timing and a Statistical Framework for Phenotype Exploration. Sleep zsac189 (2022). 10.1093/sleep/zsac189.

26. W. Truccolo, U. T. Eden, M. R. Fellows, J. P. Donoghue, E. N. Brown, A point process framework for relating neural spiking activity to spiking history, neural ensemble, and extrinsic covariate effects. J. Neurophysiol. 93, 1074–1089 (2005).

27. G. E. P. Box, D. R. Cox, An Analysis of Transformations. J. R. Stat. Soc. Ser. B Methodol. 26, 211–243 (1964).

28. Y. Benjamini, Y. Hochberg, Controlling the False Discovery Rate: A Practical and Powerful Approach to Multiple Testing. J. R. Stat. Soc. Ser. B Methodol. 57, 289– 300 (1995).

29. D. S. Falconer, Introduction to Quantitative Genetics (Pearson Education, 1996).

30. J. C. Christian, K. W. Kang, J. J. Norton, Choice of an estimate of genetic variance from twin data. Am. J. Hum. Genet. 26, 154–161 (1974).

31. J. W. Antony, et al., Sleep Spindle Refractoriness Segregates Periods of Memory Reactivation. Curr. Biol. CB 28, 1736–1743.e4 (2018).

32. T. Rusterholz, et al., Nature and Nurture: Brain Region-Specific Inheritance of Sleep Neurophysiology in Adolescence. J. Neurosci. Off. J. Soc. Neurosci. 38, 9275–9285 (2018).

