## Supplemental Figures and Tables for "Genetic Influence on the Sleep EEG is Distributed Across Multiple Traits"

**Title:**


#### Significance key

| | |
|---|---|
| 0.0000 | p < 0.001 |
| 0.0000 | p < 0.01 |
| 0.0000 | p < 0.05 |
| 0.0000 | p < 0.10 (trend) |
| 0.0000 | 95% CI lower bound > 0 (ICC table only) |

- Shading is applied to p-value cells only, except in Table 4, where each z-score cell is shaded by its matching p-value. The same thresholds are used for p-values and adjusted p-values; interpret significance by whichever column you report.
- Values are rounded to 4 decimal places; very small p-values are shown in scientific notation. "–" indicates a value that was not computed (NaN).
- The subject-level data table (data_table_best_channel.csv) is not included.

#### Supp. Table 1. Trait definitions

*Definitions of the 32 analyzed traits. Mode parameters and spectral power are computed for each of four frequency bands. Timing-pattern features are computed for sigma-high only.*

| Block | Feature | What it represents |
|---|---|---|
| **Mode parameters (each band)** | Density | How often events occur (events per minute) |
| | Frequency | Mean frequency of the events (Hz) |
| | SO power | Slow-oscillation (SO) power at the events (dB, relative to 4.5–35 Hz power) |
| | Phase Pref | SO phase of the events (radians), centered on the circular mean of all twins |
| | Duration | Duration of the events (time-frequency peaks) |
| | Prominence | Height of the events (time-frequency peaks) |
| **Timing pattern (sigma high)** | Ref period | Refractory period after an event |
| | Peak time | Time after an event when another event is most likely |
| | Peak height | Height of that peak |
| | Peak width | Width of that peak |
| **Spectral power (each band)** | Mean power | Mean spectrogram power in the band, relative to 4.5–35 Hz power (dB) |

### Supp. Table 2. Permutation z-scores for individual-trait and joint comparisons of MZ and DZ resemblance across trait groups and all 32 traits

*Permutation z-scores for the tests in Table 2 of the main text, with the same layout: z = (observed − null mean) / null SD, from 100,000 permutations of zygosity labels. Positive values mean MZ co-twins are more alike than DZ co-twins. Max Z, Mean Z and Pair Dist are the group tests (maximum Fisher z, mean Fisher z and co-twin distance) across the traits in each row, each column ("Across bands" rows) or the whole block. Cells are shaded by the matching p-value in main-text Table 2, so the two tables line up; "–" means not applicable.*

#### Mode parameters

| Group | Density | Frequency | SO power | Phase Pref | Duration | Prominence | Max Z | Mean Z | Pair Dist |
|---|---|---|---|---|---|---|---|---|---|
| Sigma high (11-16 Hz) | 1.92 | -0.55 | **2.19** | -0.28 | **2.80** | **2.56** | **2.64** | **2.76** | **2.62** |
| Sigma low (9-11 Hz) | 0.57 | 0.22 | **2.41** | -0.98 | 1.52 | 0.09 | 0.91 | 1.25 | **2.68** |
| Alpha/theta (5-9 Hz) | 1.00 | -0.70 | **2.19** | 0.57 | 1.26 | -0.96 | 1.77 | 1.29 | 2.05 |
| Delta (2-5 Hz) | 0.32 | 1.34 | 1.25 | -0.33 | 0.05 | 0.35 | 0.49 | 1.18 | **3.17** |
| Across bands: Max Z | 1.30 | 0.06 | 2.01 | -0.67 | **3.11** | 2.36 | – | – | – |
| Across bands: Mean Z | 1.83 | 0.24 | **2.20** | -0.47 | **3.04** | 1.04 | – | – | – |
| Across bands: Pair dist | **2.71** | -0.17 | **2.88** | -0.72 | **3.72** | 2.02 | – | – | – |
| **Block aggregate (24 traits)** | – | – | – | – | – | – | 1.22 | **2.86** | **3.46** |

#### Timing pattern (sigma high)

| Group | Ref period | Peak time | Peak height | Peak width | Max Z | Mean Z | Pair Dist |
|---|---|---|---|---|---|---|---|
| Sigma high (11-16 Hz) | **2.45** | 1.05 | 2.14 | 1.63 | – | – | – |
| **Block aggregate (4 traits)** | – | – | – | – | 2.26 | **3.35** | 2.07 |

#### Spectral power

| Group | Mean SpectPower | Max Z | Mean Z | Pair Dist |
|---|---|---|---|---|
| Sigma high (11-16 Hz) | 1.44 | – | – | – |
| Sigma low (9-11 Hz) | 0.16 | – | – | – |
| Alpha/theta (5-9 Hz) | 0.40 | – | – | – |
| Delta (2-5 Hz) | 0.28 | – | – | – |
| **Block aggregate (4 traits)** | – | 0.56 | 1.04 | 1.10 |

#### Aggregates by block and overall

| Group | Max Z | Mean Z | Pair Dist |
|---|---|---|---|
| Mode parameters (24 traits) | 1.22 | **2.86** | **3.46** |
| Timing pattern (4 traits) | 2.26 | **3.35** | 2.07 |
| Spectral power (4 traits) | 0.56 | 1.04 | 1.10 |
| **GRAND AGGREGATE (32 traits)** | 1.17 | **3.48** | **3.78** |

#### Supp. Table 3. Euclidean co-twin distance

*Monozygotic co-twins are closer in multivariate trait space than dizygotic co-twins. The table gives mean Euclidean distance across all traits for MZ co-twins, DZ co-twins and unrelated pairs. Significance comes from permuting zygosity labels (100,000 permutations) and testing whether DZ distance exceeds MZ distance (one-sided).*

| Statistic | Value |
|---|---:|
| Pairs (MZ) | 46 |
| Pairs (DZ) | 32 |
| Mean co-twin distance, MZ | 0.8607 |
| Mean co-twin distance, DZ | 1.1382 |
| Mean distance, strangers | 1.3762 |
| SD distance, strangers | 0.3352 |
| Stranger pairs | 12,012 |
| DZ - MZ (observed) | 0.2775 |
| Null mean | 0.0003 |
| Null SD | 0.0739 |
| z vs null | 3.7486 |
| p (one-sided) | **1.00e-04** |
| Permutations | 100,000 |

#### Supp. Table 4. Falconer estimates ($h^2$, $c^2$, $e^2$)

*Falconer estimates of additive genetic ($h^2$), shared environmental ($c^2$) and unique environmental ($e^2$) variance for each trait. Estimates outside 0-1 are shown in gray italics; they come from sampling variability in the ICCs.*

| Mode | Feature | ICC_MZ | ICC_DZ | h2 | c2 | e2 |
|---|---|---|---|---|---|---|
| Sigma high | Density | 0.6778 | 0.3050 | 0.7455 | *-0.0677* | 0.3222 |
| Sigma high | Frequency | 0.6233 | 0.7110 | *-0.1754* | 0.7987 | 0.3767 |
| Sigma high | SO power | 0.5482 | -0.1041 | *1.3046* | *-0.7564* | 0.4518 |
| Sigma high | Phase Pref | 0.0743 | 0.1564 | *-0.1644* | 0.2386 | 0.9257 |
| Sigma high | Duration | 0.8218 | 0.3052 | *1.0331* | *-0.2113* | 0.1782 |
| Sigma high | Prominence | 0.8147 | 0.3818 | 0.8657 | *-0.0510* | 0.1853 |
| Sigma high | Mean power | 0.7067 | 0.4728 | 0.4678 | 0.2389 | 0.2933 |
| Sigma high | Ref period | 0.6783 | 0.1629 | *1.0308* | *-0.3525* | 0.3217 |
| Sigma high | Peak time | 0.4656 | 0.2406 | 0.4499 | 0.0156 | 0.5344 |
| Sigma high | Peak height | 0.6717 | 0.2459 | 0.8516 | *-0.1799* | 0.3283 |
| Sigma high | Peak width | 0.3126 | -0.0592 | 0.7435 | *-0.4309* | 0.6874 |
| Sigma low | Density | 0.7443 | 0.6655 | 0.1576 | 0.5867 | 0.2557 |
| Sigma low | Frequency | 0.5343 | 0.4428 | 0.1830 | 0.3513 | 0.4657 |
| Sigma low | SO power | 0.5569 | -0.0676 | *1.2489* | *-0.6920* | 0.4431 |
| Sigma low | Phase Pref | 0.3642 | 0.5382 | *-0.3480* | 0.7122 | 0.6358 |
| Sigma low | Duration | 0.8523 | 0.7098 | 0.2851 | 0.5672 | 0.1477 |
| Sigma low | Prominence | 0.4224 | 0.4085 | 0.0278 | 0.3946 | 0.5776 |
| Sigma low | Mean power | 0.7716 | 0.7532 | 0.0369 | 0.7347 | 0.2284 |
| Alpha/theta | Density | 0.7529 | 0.6289 | 0.2481 | 0.5048 | 0.2471 |
| Alpha/theta | Frequency | 0.4802 | 0.6315 | *-0.3027* | 0.7829 | 0.5198 |
| Alpha/theta | SO power | 0.5706 | 0.0169 | *1.1074* | *-0.5368* | 0.4294 |
| Alpha/theta | Phase Pref | 0.3527 | 0.2283 | 0.2489 | 0.1038 | 0.6473 |
| Alpha/theta | Duration | 0.7906 | 0.6067 | 0.3677 | 0.4229 | 0.2094 |
| Alpha/theta | Prominence | 0.5219 | 0.6706 | *-0.2973* | 0.8192 | 0.4781 |
| Alpha/theta | Mean power | 0.7160 | 0.6667 | 0.0986 | 0.6175 | 0.2840 |
| Delta | Density | 0.5760 | 0.5152 | 0.1216 | 0.4544 | 0.4240 |
| Delta | Frequency | 0.6186 | 0.2941 | 0.6491 | *-0.0305* | 0.3814 |
| Delta | SO power | 0.3378 | -0.0212 | 0.7180 | *-0.3802* | 0.6622 |
| Delta | Phase Pref | 0.2585 | 0.3571 | *-0.1973* | 0.4558 | 0.7415 |
| Delta | Duration | 0.3672 | 0.3588 | 0.0167 | 0.3505 | 0.6328 |
| Delta | Prominence | 0.3819 | 0.3098 | 0.1442 | 0.2377 | 0.6181 |
| Delta | Mean power | 0.7157 | 0.6750 | 0.0813 | 0.6343 | 0.2843 |
| **Mean of 32 traits** | | **0.5650** | **0.3815** | **0.3671** | **0.1979** | **0.4350** |

#### Supp. Figure 1. Correlations between the 32 traits

*Pearson correlations between all 32 traits across all participants, with traits grouped by frequency band. Red is a positive correlation and blue is a negative one.*

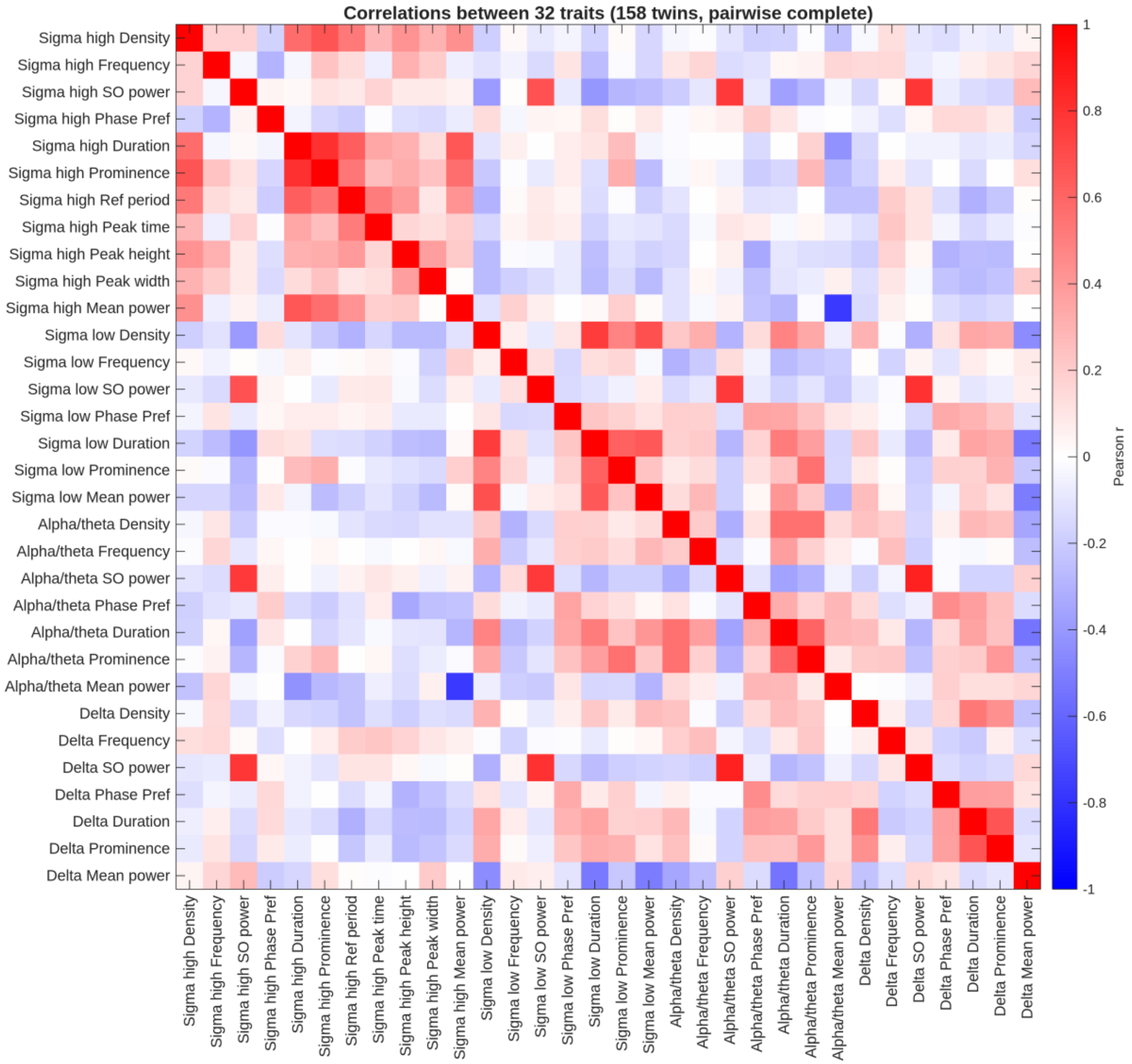
